# Phen2Disease: A Phenotype-driven Semantic Similarity-based Integrated Model for Disease and Gene Prioritization

**DOI:** 10.1101/2022.12.02.518845

**Authors:** Weiqi Zhai, Xiaodi Huang, Nan Shen, Shanfeng Zhu

## Abstract

By utilizing the Human Phenotype Ontology (HPO), recent approaches to prioritizing disease-causing genes for patients become popular. However, these approaches do not comprehensively use information about phenotypes of diseases and patients. We present a new method called Phen2Disease that calculates similarity scores between two phenotype sets of patients and diseases by which to prioritize diseases and genes. Specifically, we calculate three scores of information content-based similarities using the phenotypes, and their combination as the respective benchmarks, and integrate them as a final score. Comprehensive experiments were conducted on six real data cohorts with 2051 cases and two simulated data cohorts with 1000 cases. Compared with the three state-of-the-art methods, if we only use phenotype information and HPO knowledge base, Phen2Disease outperformed all of them, particularly in cohorts with the less average numbers of HPO terms. We have found that patients with higher information content scores had more specific information so their predictions would be more accurate. In addition, Phen2Disease has high interpretability with ranked diseases and patient HPO terms provided.

## INTRODUCTION

In the clinical field, physicians need to diagnose diseases and causative genes based on the observed clinical features of patients. Thousands of genes are known to cause Mendelian diseases, and hundreds of new disease-causing genes are discovered every year [1–2]. It is difficult and time-consuming for physicians to make diagnoses, especially in the field of rare genetic diseases. According to the preliminary statistics, it takes a few months or even years for specialists to identify a patient’s causative genes [3–4]. The current standard method for the timely and accurate diagnosis of Mendelian diseases is exome sequencing. It can detect candidate deleterious gene variants in protein-coding regions at a low cost, significantly improving the effective identification of disease genes [5–8]. However, dozens or hundreds of such deleterious gene variants can be predicted. Therefore, the most likely gene variants need to be ranked for further screening.

For this purpose, phenotype-driven screening-based approaches are suitable for filtering out many disease-causing genes. In fact, phenotype-driven prioritization of candidate genes and diseases is becoming a well-established approach to genomic diagnostics in rare diseases [9–18]. Essentially, phenotypic approaches screen for candidate diseases by searching for those with similar observed phenotypic abnormalities that are associated with carried disease [19]. The application of phenotypic methods necessitates the recording of clinical characteristics or phenotypic information. Many databases storing disease phenotypic data are currently available. Examples of the most widely used ones are the Human Phenotype Ontology (HPO) database [20–22], OMIM [23], and Orphanet [24]. HPO contains a comprehensive description of human phenotypes organized as a hierarchical rooted directed acyclic graph (DAG). For version 2021.04.13, HPO provides such rooted DAGs for 15872 different phenotypes, with Human Phenotype Ontology Annotation (HPO-A) consisting of 4555 genes, 7839 diseases, and 8436 direct associations between HPO phenotypes and diseases. With these available databases, it is possible that phenotypic approaches automatically rank candidate genes by using phenotypic information in the databases. Physicians then focus their attention on the most likely genes at the top of the ranked list. In this way, the scope of the diagnosis can be significantly narrowed so that the diagnosis accuracy can be improved with reduced time.

Currently, many tools are available that use phenotypic methods to automatically rank candidate genes based on given phenotypes [25–30]. These tools can roughly be classified into two categories. The first type of phenotypic analysis approach prioritizes the diseases first, followed by the prioritization of candidate causative genes. As shown in Figure 1, this type of approach uses the associations among phenotypes, diseases and disease genes to build a knowledge base that allows the prioritization of candidate diseases and genes. Most recent examples include LIRICAL [25] and Phrank [26]. These approaches differ mainly in how they match patient phenotypes with disease phenotypes in terms of their corresponding HPO terms in the hierarchical HPO DAG. Specifically, by using the likelihood ratio (LR) for interpretable clinical genomics, LIRICAL calculates the ratio between two conditional probabilities that phenotypes are observed with and without a given disease. It further integrates the observed genotypes to predict pathogenicity. As such, clinicians can use LIRICAL to assess the contribution of each phenotypic abnormality to a diagnosis. However, LIRICAL has a strong assumption that the phenotypes of a patient are mutually independent. If an HPO term of a patient phenotype cannot be matched with that of the target disease phenotypes, LIRICAL uses its related kin terms on the HPO tree instead to calculate the likelihood ratio with a heuristic weighting scheme. In contrast, Phrank calculates similarity scores using the information content of overlapping patient and disease phenotypes sets. Phrank has the following drawbacks. First, it uses only local information, i.e., the information of parent and sibling phenotypes, to calculate the information content of phenotype nodes. Second, it does not distinguish the importance of nodes at different levels in an HPO ontology. Finally, Phrank considers only phenotypes in the intersection of the sets of patient and disease phenotypes, ignoring all others in the two sets.

**Figure 1.**
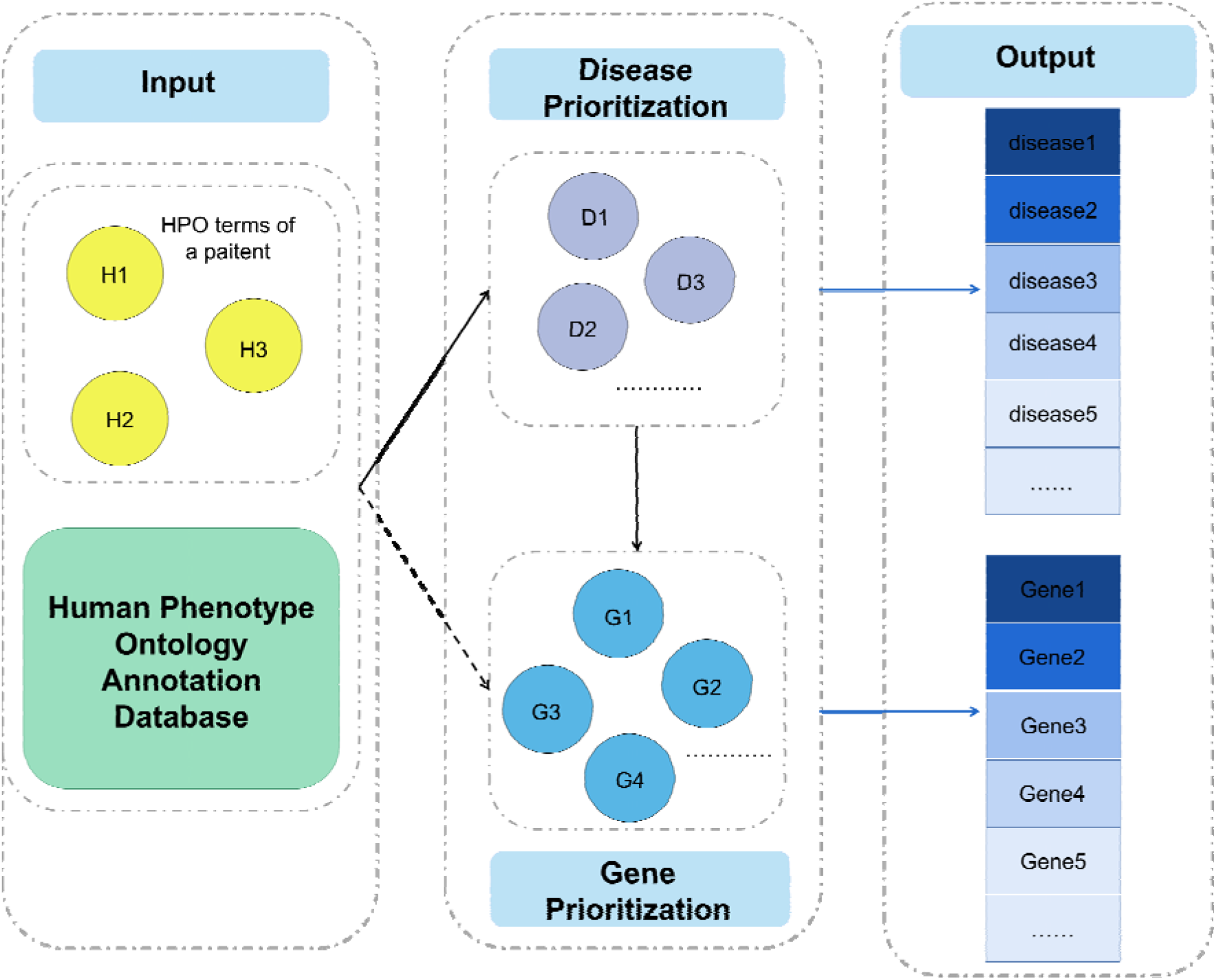
Diagnosing diseases and disease-causing genes of patients. Prioritization of candidate diseases uses patient HPO terms and Human Phenotype Ontology Annotation Database. Considering the associations between diseases and genes, prioritization of candidate genes could be obtained through prioritization of candidate diseases. The solid lines represent direct associations and the dashed line represents an indirect association.

The second category of phenotype methods also aims to find relationships between phenotypes and candidate genes for prioritizing candidate genes, but without identifying the disease first. Typical examples of the second category include Phenolyzer [27], Phen2Gene [28], AMELIE [29], and PhenoApt [30]. Compared with these methods, the first category of approaches provides good interpretability, which is the focus of our work. In conclusion, all these methods for candidate disease and gene prioritization could be improved. New phenotype-driven disease and gene prioritization approaches are urgently needed.

An effective phenotype-based approach should make full use of all the information about the phenotypes of a patient and a disease and their matched HPO terms. However, this faces the following challenges. First, the observed patient phenotypes often include incorrect, incomplete, and noisy phenotypes, which may miss important symptoms. The phenotypes of a disease may be not specific, covering the general symptoms of most patients. And patients with the same disease may have several subtypes and thus have various phenotypes [31]. Second, there are different ways to extract standardized terms such as HPO from clinical notes. Further, different physicians may use different phenotype terms even if they refer to the same phenotypes. Third, the HPO terms associated with phenotypes of a patient and a disease are not always matched exactly with those on the HPO ontology. Therefore, we should develop a new way of accurately matching the phenotypes between a disease and a patient.

In this work, we introduce a new method for ranking diseases and genes called Phen2Disease that prioritizes the relationships between patient phenotypes and diseases. Fully considering the graph structural information of HPO, we use the information content to measure the importance of each HPO term. First, patient-centric and disease-centric similarity scores between two phenotype sets are calculated by using phenotypes of a patient and disease as a respective benchmark. Then, we calculate patient-disease-centric similarity scores using both patient and disease phenotypes as benchmarks. Using comprehensive information, we finally integrate the patient-centric score and patient-disease-centric score to produce the final similarity scores between the two sets of phenotypes, by which the candidate diseases and genes can be prioritized. Phen2Disease has demonstrated excellent performance for disease and gene prioritization in six real cohorts, with good clinical interpretability. Compared with the three state-of-the-art methods (Phrank, PhenoApt and LIRICAL), if we only use phenotype information and HPO knowledge base, Phen2Disease outperformed all of them, particularly in cohorts with the less average numbers of HPO terms. We have found that patients with higher information content scores had more specific information so their predictions would be more accurate. Additionally, the superior performance of Phen2Disease has been validated on two simulated cohorts with added noise.

## MATERIAL AND METHODS

### Calculating a Phen2Disease score between two sets of phenotypes

For a given patient, Phen2Disease takes as input a list of the phenotypes and another list of candidate diseases and genes. Based on a user preference, Phen2Disease then outputs a ranking score for each candidate disease that may be caused by one or more genes on the patient candidate gene list selected by users or provided based on the available knowledgebase. The higher the score, the higher likelihood of the corresponding disease that can explain patient phenotypes in a given set.

The overall procedure for Phen2Disease is illustrated in Figure 2. Given a set of patients’ (or disease) phenotypes, we first propagate the phenotypes (HPO terms) along the HPO ontology, then filter out HPO terms that have descendants in the set, and finally obtain a reduced set of phenotypes. Given a set of patient phenotypes *P* and another set of disease phenotypes *D*, for two phenotype terms *t* ∈ *P* and *t*’∈ *D*, Phen2Disease starts by calculating the information content (IC) of each HPO term *t*. Specifically, the IC of an HPO term *t* is defined as follows:

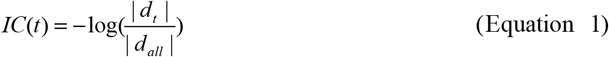

where |*d_t_*| is the number of diseases associated with the HPO term *t* in the HPO database, and |*d_all_*| is the number of all diseases in the HPO database. Second, we calculate the similarity score between any two HPO terms *t* and *t*’ using the Lin measure [32]:

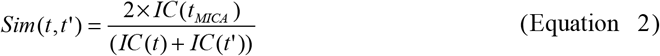

where MICA represents the lowest common ancestor of the two terms.

**Figure 2.**
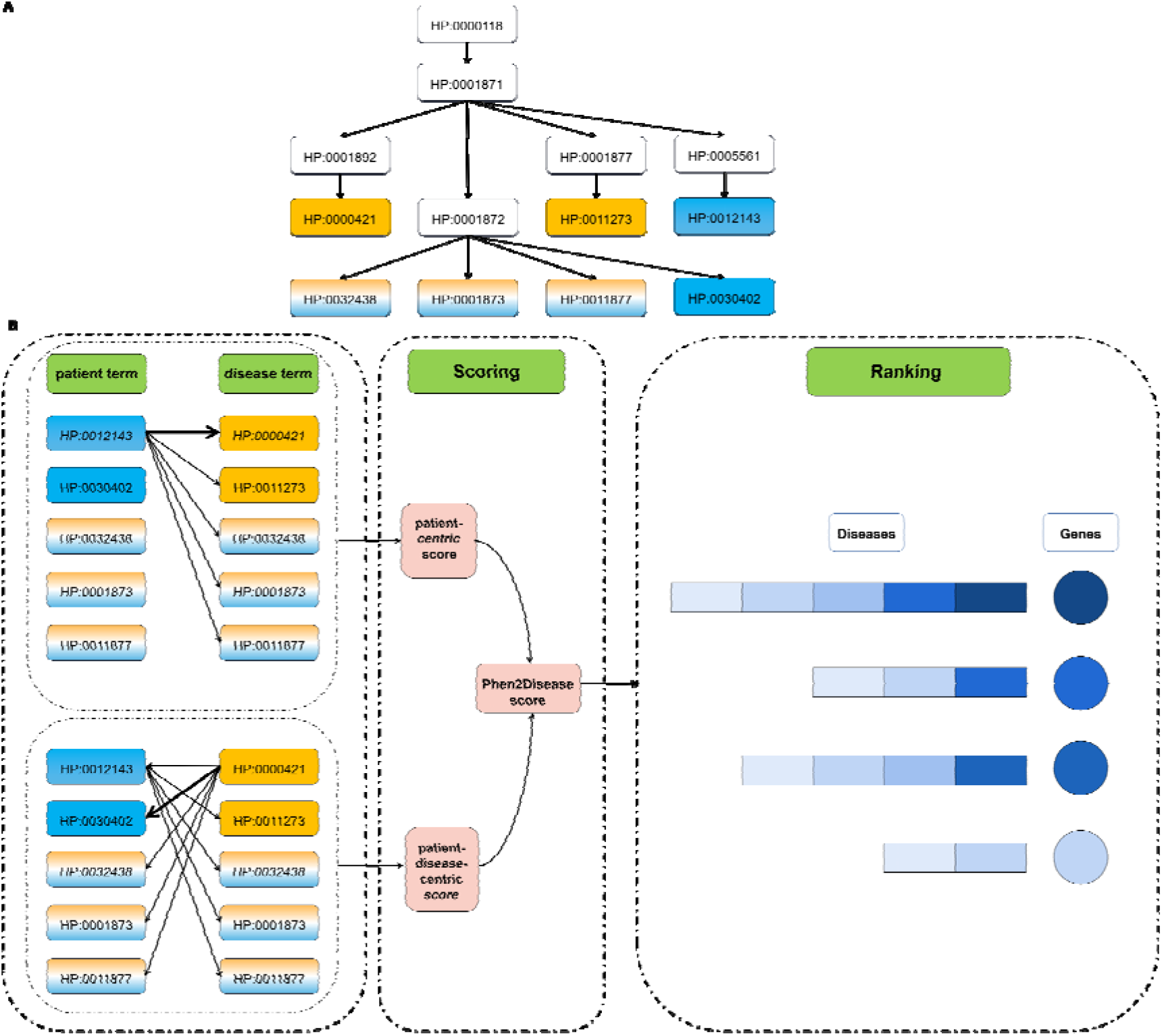
The overview of Phen2Disease. **A.** Annotation of patient phenotypes with corresponding disease phenotypes on the HPO. Overlapping phenotype terms are highlighted in the gradient color of blue and yellow, with patient-specific phenotypes in blue and disease-specific phenotypes in yellow. **B.** Using Information Content, we calculate the patient-centric similarity scores (Equation 3) between patient phenotypes and candidate disease phenotypes. Then we obtain a patient-disease-centric similarity score (Equation 5) by considering both patient and disease as benchmarks. Finally, we integrate the patient-centric score and the patient-disease-centric score to produce the final similarity scores of the patient for each candidate disease (Equation 6). In this way, the candidate diseases of a patient are ranked in terms of their corresponding scores (The thick line represents the maximum score of matched phenotypes). A candidate gene could be associated with multiple diseases, so the Phen2Disease score for a gene is defined as the maximum score corresponding to the most likely disease caused by this particular gene.

After calculating the similarity between two HPO terms, we can calculate the similarity scores between patient phenotypes in one set and the disease phenotypes in another set by using the human phenotype ontology. Specifically, for a set of patient phenotypes, we first compute a sum of the maximum similarity scores between each term in a set of patient phenotypes and each term in a set of disease phenotypes, weighted by the IC score of this particular term. We then normalize a patient-centric similarity score by the sum of the weights (IC of each term) as:

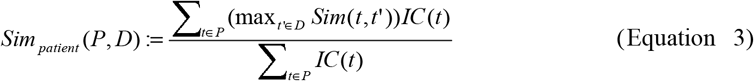

For the disease-centric similarity score, we calculate the similarity score between each term in a set of disease phenotypes and each term in a set of patient phenotypes, which is similar to calculating the patient-centric similarity:

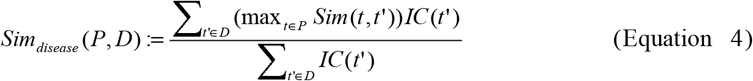

Both of these methods can calculate a patient’s candidate disease score. However, *Sim_patient_* and *Sim_disease_* emphasize patient-centric or disease-centric information, respectively. Inspired by our previous work on computing semantic similarity between two documents based on Medical Subject Headings (MeSH) [33], we make use of the term information from both sets of disease and patient phenotypes to produce patient-disease-centric similarity scores as follows:

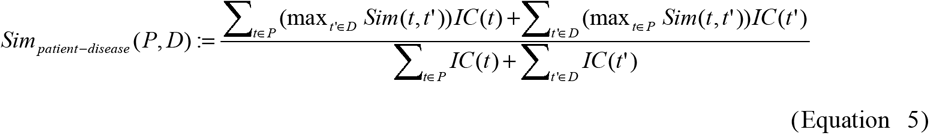

From the clinical point of view, a process of calculating patient-centric scores matches well with a diagnostic process because it is a common practice to start a diagnosis by examining the phenotypic characteristics of a patient. It is also beneficial to use accurate disease phenotype information from a disease perspective. In reality, a diagnosis is always accompanied by noise information from different resources, which often are not consistent with each other. Using the complete information about patients and diseases, we integrate *Sim_patient–disease_* and *Sim_patient_* for an effective clinical diagnosis. Finally, the Phen2Disease score between the sets of phenotypes of patients and diseases is calculated as:

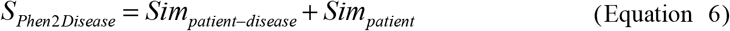

To differentiate the Phen2Disease variants. we denote *Sim_patient_* as “Phen2Disease-patient”, *Sim_disease_* as “Phen2Disease-disease”, and *sim_patient–disease_* as “ Phen2Disease-double” in the following.

### The patient-specific Phen2Disease score of a candidate gene

Given the above definition of the Phen2Disease score, the patient-specific Phen2Disease score of disease is simply the similarity score between a set of disease phenotypes associated with those in the knowledgebase and an input set of patient phenotypes. By using information about patient-disease-gene rather than about patient-disease, the Phen2Disease score of a candidate gene is defined as the maximal Phen2Disease score for a disease that is most likely caused by this particular gene, according to the knowledgebase (e.g. HPO).

### The baseline methods

Besides Phen2Disease, we present the BASE_IC approach that directly quantifies the degree of the relationship based on the common terms in a set of patient phenotypes and a set of candidate genes or disease phenotypes. Specifically, BASE_IC calculates the similarity score of a disease or gene for a patient by first finding common terms in both sets of phenotypes and then summing their corresponding information content scores:

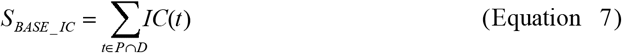

Note that both BASE_IC and Phrank calculate the sum of information content scores for the intersection of two sets of phenotypes, but BASE_IC uses the global information content. In our experiments, we compare Phen2Disease with the baseline methods of Phenolyzer, Phen2Gene, Phrank, PhenoApt, LIRICAL and AMELIE, including BASE_IC. Among them, AMELIE [29] differs from other methods in that it augments the knowledge base by mining biomedical literature. All these baseline methods can be used to rank candidate genes. In particular, Phrank, PhenoApt and LIRICAL can also be used to rank candidate diseases. As given in Table 1, we compare the phenotypic information utilized by these methods.

**Table 1.** Comparisons of phenotypic information used by the different methods. In this table, “patient-” stands for patient-centric, “LR” for likelihood ratio, “MD” for matrix decomposition, “Intersection” for the intersection of patient phenotypes and disease phenotypes, and “All” for all phenotypes of patient and disease.

|  | LIRICAL | Phrank | BASE_IC | Phen2Disease | PhenoApt |
| --- | --- | --- | --- | --- | --- |
| Model/<br>Method | LR | IC | IC | IC | MD |
| Phenotype<br>utilization | patient | Intersection | Intersection | All | All |
| Similarity<br>between<br>phenotypes | Ad-Hoc | Binary | Binary | Lin [32] | Dot product<br>with<br>user-specifi<br>ed weights |

### Evaluation datasets of true diagnostically validated patients

To validate the performance of Phen2Disease, we conducted experiments by comparing it with the baseline methods against the six cohorts. As summarized in Table 2, we describe the six cohorts as follows.

**Table 2.**
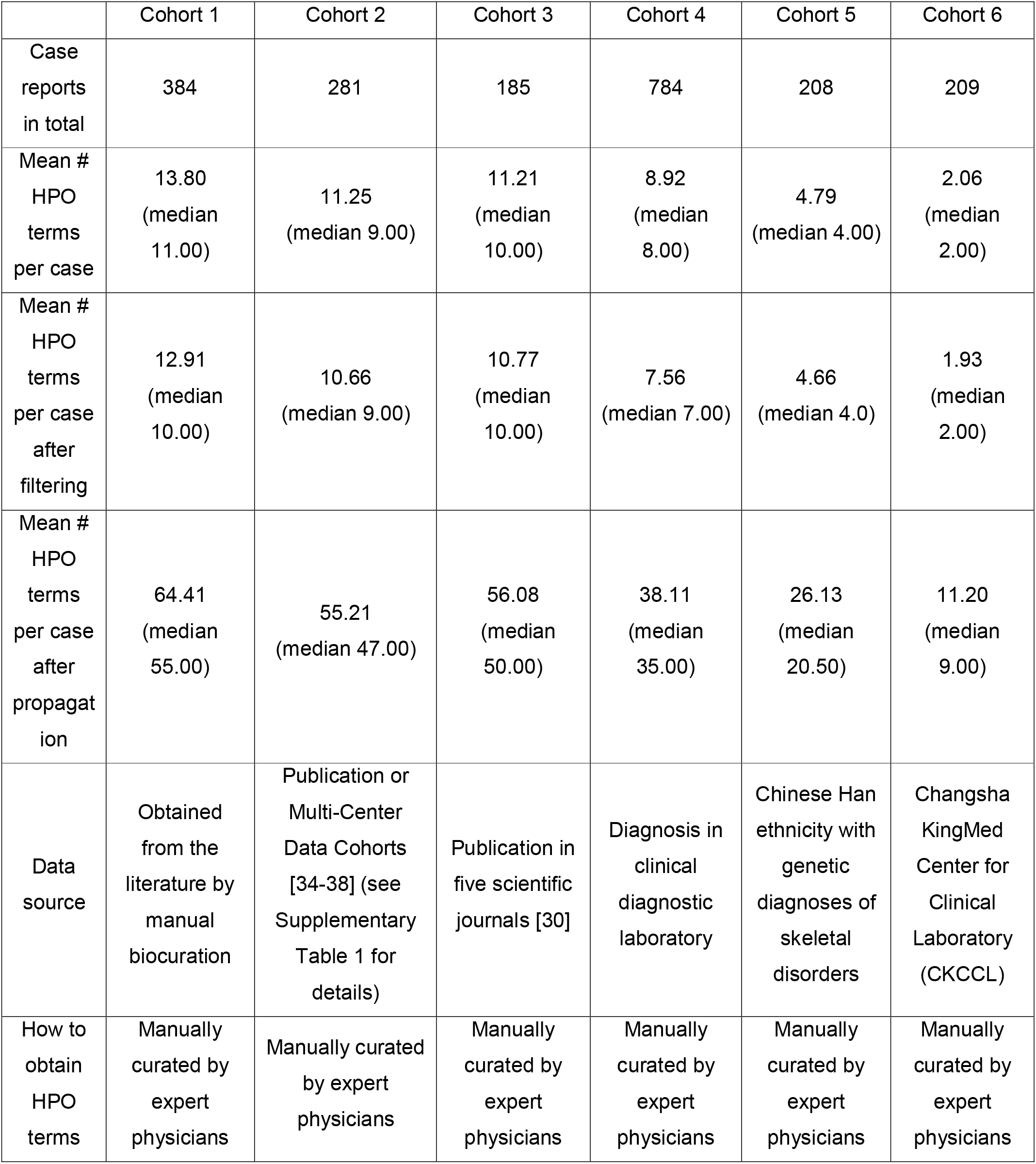
Statistics of the datasets used in the experiments. According to the true path rule: if an entity is annotated as a possible term, it is implicitly annotated by all ancestor terms of that term. The “propagation” means that all ancestor nodes of each phenotype in the phenotype set are considered. The “filtering” means that if there are ancestor-descendant node relationships in the phenotype collection, only the descendant nodes are retained.

The real data cohorts we used are all from the data cohorts used in the compared methods, except for the articles of Phrank and Phenolyzer for which no data cohorts are publicly available. For Cohort 1, we used the same data cohort as Robinson et al. [25] – 384 cases, which were derived from the literature by manual biocuration, with each describing a single published case report. In our method, we focused only on the phenotypic information of the patients. For Cohort 2, we used the same data cohort as Zhao et al. [28]. Cohort 2 consists of 281 patients, with each phenotype manually curated by the doctors. The cases came from five types of resources [34–38], such as the American Journal of Human Genetics Article, Columbia University Medical Center, and the Department of Genomic Diagnostics at the Children’s Hospital of Philadelphia. For Cohorts 3-5, we used the same data cohort as Chen et al. [30]. The resources of Cohort 3 were from the publications in five scientific journals (American Journal of Medical Genetics Part A, BMC Medical Genetics, BMC Pediatrics, Frontiers in Genetics, or Molecular Genetics & Genomic Medicine). Cohort 4 consists of 784 singleton-affected individuals with molecular diagnoses from a clinical diagnostic laboratory. The clinical diagnostic criteria were under the American College of Medical Genetics/Association for Molecular Pathology (ACMG/AMP) guidelines [39–41]. Cohort 5 consists of 208 individuals of Chinese Han ethnicity with genetic diagnoses of skeletal disorders from 2009 to 2019, as part of the Deciphering disorders Involving Scoliosis and COmorbidities (DISCO) study group [42–44]. For Cohort 6, we used the same data cohort as Yuan et al. [45]. In particular, Cohort 6 consists of 209 in-house cases from Changsha KingMed Center for Clinical Laboratory (CKCCL). Genetic analysts screened and collated all HPO terms for these cases by removing ambiguous terms. As a result, the average number of HPO terms per case is 2.06 in the experiments.

### Evaluation datasets on simulated patients

To further test the robustness of our model under large datasets with noise, we built two simulated datasets with a sample size of 1000 and added noises. Firstly, we simulated 1000 patients with their diseases from 7839 diseases in the HPO database. The phenotypes of each patient were generated from the distribution of phenotypic data corresponding to real diseases. Comparing the HPO term sets of 384 real patients (Cohort 1) with their corresponding diseases, we first calculated the ratio of the total number of terms in the phenotypes of real patients to that of terms in their corresponding diseases and then matched their relationships of the term nodes on HPO. The proportions of the same nodes, parent nodes, child nodes, ancestor nodes, descendant nodes, sibling nodes, and other possible nodes were finally calculated for all real patients. The data distribution of patients compared to real diseases was obtained by averaging these proportions. The distribution of the resulting data is shown in Table 3.

**Table 3.**
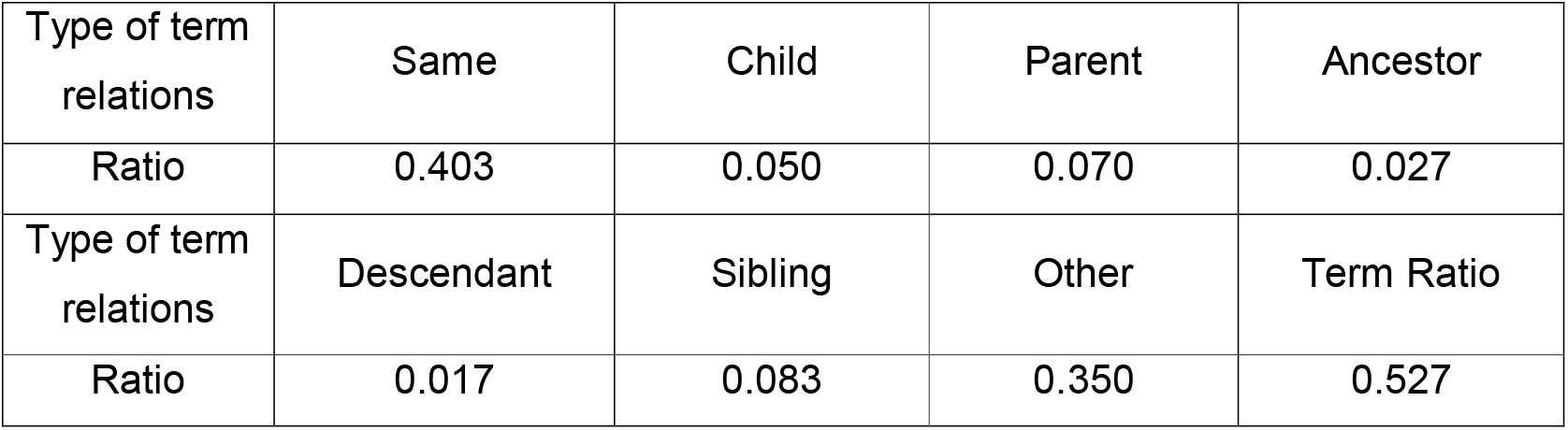
Data distribution of patients compared to real diseases. The term “Ratio” means the average ratio of the number of HPO terms in P to the number of HPO terms in D; “Same” means the term t is Identical to One of the Terms to Which phenotypes set D Is Annotated; “Child” means t Is a Child Term of One or More of the Terms to Which D Is Annotated; “Ancestor” means t Is an Ancestor of One or More of the Terms to Which D Is Annotated; “Descendant” means t Is a Descendant of One or More of the Terms to Which D Is Annotated; “Sibling” means t and Some Term to Which D Is Annotated Have a Non-root Common Ancestor, and are of the same depth; “Relatives” means t and Some Term to Which D Is Annotated Have a Non-root Common Ancestor and are not of the same depth.

| Type of term relations | Same | Child | Parent | Ancestor |
| --- | --- | --- | --- | --- |
| Ratio | 0.403 | 0.050 | 0.070 | 0.027 |
| Type of term relations | Descendant | Sibling | Other | Term Ratio |
| Ratio | 0.017 | 0.083 | 0.350 | 0.527 |

We simulated the second cohort by following the method of Chen et al. [30]. Each disease-causing gene has its corresponding directly related phenotypes. First, two-thirds of directly related phenotypes were randomly removed. Second, the remaining half of directly related phenotypes were replaced with indirectly related phenotypes (i.e., the ancestors of the directly associated phenotypes). Third, unassociated phenotypes were randomly generated and added. Finally, each case in the test set of simulated patients contains 25% of directly related phenotypes, 25% of indirectly related phenotypes, and 50% of unrelated phenotypes. The simulated test set contains 1000 cases in total.

### Performance evaluations

For the experiments, we used the Python implementation of Phenolyzer, Phen2Gene, Phrank, PhenoApt, and LIRICAL, respectively. The procedure of the experiments is as follows. For each case, we input a list of HPO terms into the tool. Each case was tested with its causative gene included in the list of resultant candidate genes. For AMELIE, we conducted the experiments on its website version by saving the diagnosis file for each case and locating the causative gene in the candidate gene list provided by AMELIE.

We compared our method with the baseline methods by examining their ranked causative genes. The ranking result from each method is “top 1”, “top 5”, “top 10”, “top 20”, “top 50”, “top 100”, or “other”. Note that if a case had no causative gene in the returned candidate gene list, it was categorized as “other”. Also, we used mean reciprocal rank (MRR) as the metric to evaluate the performance of all compared methods. It is worth noting that we did not evaluate the MRR metrics of PhenoApt, since PhenoApt outputs only the top 100 candidate genes (or diseases), without producing a full gene (or disease) ranking. Furthermore, we use the Wilcoxon signed rank test to compare the differences in the prioritization ability of Phen2Disease and all competing methods. For fairly comparing methods that determine the likelihood of a caused gene, we selected all genes and diseases recorded in the HPO database as candidate genes and diseases.

## RESULTS

Phen2Disease takes a set of phenotypes for each patient as input for candidate genes ranking. We evaluated the performance of Phen2Disease in terms of its ability to prioritize disease-causing genes from a list of candidate genes. In the first part of the experiments, we compared the performance of Phen2Disease with other compared methods against the real independent cohorts. In the second part of the experiments, we used cohorts of simulated patients instead to evaluate their performance.

### Phen2Disease performed well on six real-world, multi-cohorts

Among our real data cohorts, only Cohort 1 (N=384) disease had annotations for each case. So, we used data Cohort 1 to test the performance of Phen2Disease on ranking candidate diseases by comparing with BASE_IC, LIRICAL, PhenoApt, and Phrank. As shown in Supplementary Figures 1-2, Phen2Disease outperformed all other compared methods. In particular, the diseases Phen2Disease ranked first are the true ones in 28.91% of cases, with those correctly ranked first by BASE_IC, LIRICAL, Phrank, and PhenoApt being 25.52%, 24.48%, 21.61%, and 21.61%, respectively. The mean reciprocal rank (MRR) of Phen2Disease in Cohort 1 for ranking diseases was 38.64, while that of BASE_IC, LIRICAL, and Phrank was 35.31, 32.72, and 31.19, respectively (see Supplementary Table 2). The performance of our Phen2Disease on ranking diseases is much better than that of all the compared methods, which indicates that Phen2Disease has a strong diagnostic power in the preferential selection of diseases.

Compared to the methods of BASE_IC, Phrank, LIRICAL, and PhenoApt that rely only on HPO knowledgebase for gene prioritization, our model Phen2Disease has achieved the first in terms of MRR on all six cohorts of data. By using the Wilcoxon signed rank test, all the results from Phen2Disease were statistically significantly (p<0.01) improved over those from BASE_IC on all the cohorts, those from Phrank on two data cohorts, and those from LIRICAL on three data cohorts. Besides, Phen2Disease produced similar priority ranking results as AMELIE (see Supplementary Table 3), though AMELIE uses the extended knowledge base beyond HPO. Specifically, for candidate gene prioritization, Phen2Disease performed best in the one independent, real-world cohort (Cohort 5), first (tied) in the three cohorts (Cohort 1, Cohort 4 and Cohort 6), and second place in another two cohorts (Cohort 2 and Cohort 3), in terms of the TopK (K=1) metric (see Supplementary Tables 4-14). Take Cohort 5 as an example (Figure 3E). Phen2Disease performed well and ranked the true causative gene first in 24.52% of cases. In contrast, PhenoApt, Phrank, BASE_IC, Phen2Gene, Phenolyzer, AMELIE and LIRICAL ranked 13.94%, 10.58%, 8.65%, 8.17%, 8.17%, 5.77%, and 3.85%, respectively. The mean reciprocal rank (MRR) of Phen2Disease in Cohort 5 was 31.79, while the MRR of Phrank, BASE_IC, Phen2Gene, Phenolyzer, AMELIE and LIRICAL was 19.54, 14.86, 13.43, 15.06, 15.47, and 10.27, respectively (see Supplementary Table 13). Further, we found that as a real data cohort with some noise, Cohort 5 has fewer mean HPO terms (N=4.66) under a large disease category like skeletal disorders. Our model showed a good ability to prioritize disease-causing genes for patients with skeletal diseases in clinical trials.

**Figure 3.**
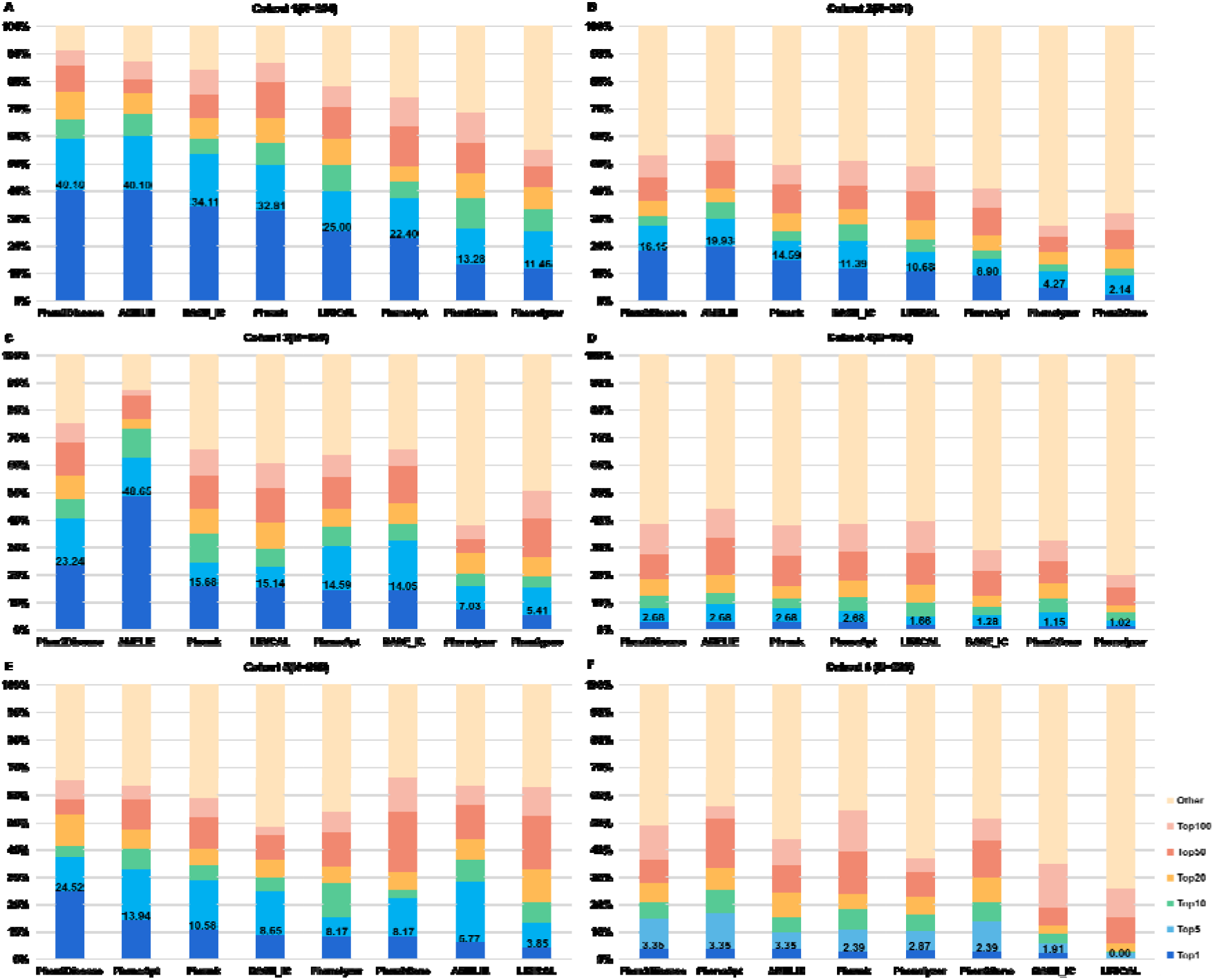
Comparisons of Gene ranking results on the six real cohorts. **A.** Comparisons of the eight methods of Phen2disease, BASE_IC, Phrank, Phenolyzer, Phen2Gene, LIRICAL, PhenoApt, and AMELIE for gene prioritization on Cohort 1 (384 cases). **B.** Comparisons on Cohort 2 (281 cases). **C.** Comparisons on Cohort 3 (185 cases). D. Comparisons on Cohort 4 (784 cases). E. Comparisons on Cohort 5 (208 cases). F. Comparisons on Cohort 6 (209 cases).

It is worth mentioning that AMELIE performs much better than all other compared methods in Cohort 3; however, it performs poorly in Cohorts 5 and 6. For this reason, we analyzed the results by carefully checking 185 cases in Cohort 3. From the published scientific journals, all these cases could be found in PubMed’s data repository for corresponding case reports. Besides using all 29 million PubMed abstracts, AMELIE analyzed hundreds of thousands of full-text articles to find information supporting the causality and associated phenotypes of most published genetic variants. Therefore, the gene prioritization effect of AMELIE is far superior to other methods (see Supplementary Table 11). In contrast, Cohorts 5 and 6 have only 4.66 and 1.93 HPO terms per patient on average, respectively. If a case has too few HPO terms, AMELIE assigns the same score to a large number of candidate genes, thus losing its ability to prioritize them. In fact, only 5.77% and 3.35% of the correct candidate genes were ranked first by AMELIE (see Supplementary Tables 13-14), respectively, which shows its shortcomings.

### Phen2Disease greatly improves the accuracy of gene and disease prioritization in simulated datasets

For the first set of our simulated data, we reported the disease prioritization of data Cohort 1 (Figure 4A). Phen2Disease ranked the true disease first in 61.20% of the cases, with PhenoApt, BASE_IC, LIRICAL and Phrank as 34.00%, 29.80%, 29.10%, and 22.90%, respectively. The mean reciprocal rank (MRR) of Phen2Disease in Cohort 1 for disease ranking was 70.66, followed by BASE_IC (38.28), LIRICAL (40.42), and Phrank (32.89), respectively (see Supplementary Table 15). Furthermore, we performed Wilcoxon signed rank test for these results. The p-value of the test with BASE_IC, LIRICAL, and Phrank was 1.7*10^-61^, 2.4*10^-65^, and 8.5*10^-69^, respectively (see Supplementary Table 3).

**Figure 4.**
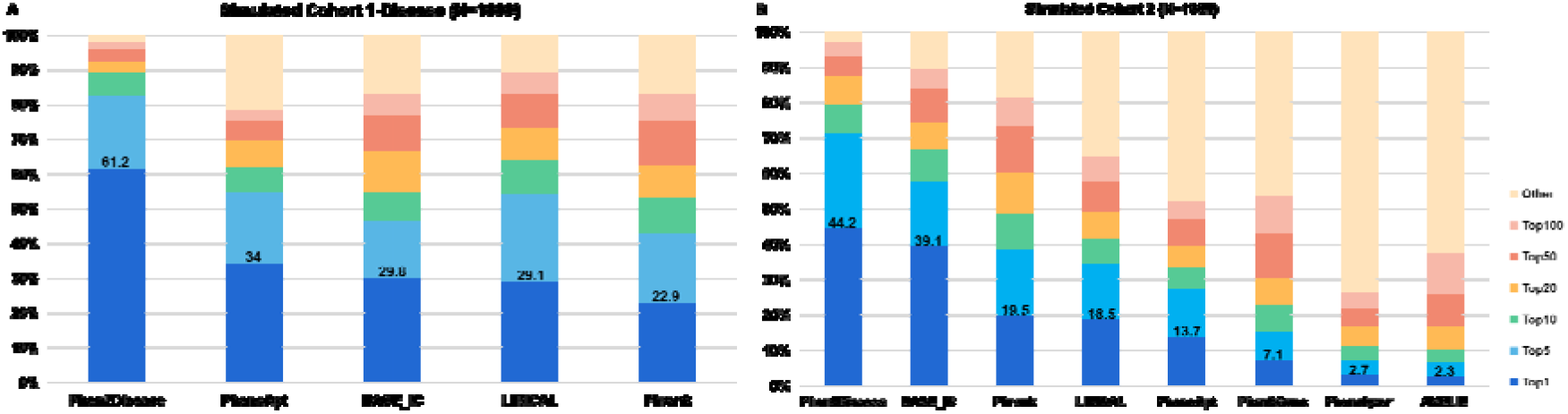
Gene and disease ranking results in two simulated cohorts. **A.** Comparisons of the five methods of Phen2disease, BASE_IC, Phrank, LIRICAL, and PhenoApt for disease prioritization in the simulated Cohort 1 (1000 cases). **B.** Comparisons of the eight methods for gene prioritization in the simulated Cohort 2 (1000 cases).

For the second set of simulated data, we followed the method of Chen et al. [29] and generated 1000 simulated cases (Figure 4B). Phen2Disease ranked first in 44.20% of cases, and BASE_IC, Phrank, LIRICAL, PhenoApt, Phen2Gene, Phenolyzer, and AMELIE ranked 39.10%, 19.50%, 18.50%, 13.70%, 7.10%, 2.70%, and 2.30%, respectively. The mean reciprocal rank (MRR) of Phen2Disease in simulated Cohort 2 was 56.12, with BASE_IC, Phrank, LIRICAL, Phen2Gene, Phenolyzer, and AMELIE being 48.21, 28.87, 26.33, 11.92, 5.57, and 5.53, respectively (Supplementary Table 16). Wilcoxon signed rank test confirmed the superiority of Phen2Disease (see Supplementary Table 3 for the detailed information). We can find that these cases in simulated Cohort 2 are not reported in the literature before and thereby AMELIE loses the advantage.

### Phen2Disease outperforms its three variants

The Phen2Disease can be regarded as an integrated model with different components. As such, we conducted experiments to test the performance of the components of Phen2Disease in all data cohorts.

First, we found that Phen2Disease performed best in all cohorts. Phen2Disease-patient and Phen2Disease-double performed well in both the real and simulated data cohorts. While Phen2Disease-disease performs the worst in all data cohorts. For example, for Cohort 1 (N=384), Phen2Disease-patient and Phen2Disease-double ranked 34.64% and 26.56% of the real disease-causing genes in Top1, respectively, but only 2.86% for patient-disease. Phen2Disease-patient and Phen2Disease-double ranked 25.00% and 21.09% of the real diseases in Top1, respectively, but only 1.56% for Phen2Disease-disease. The results of disease prioritization for Cohort 1 are shown in Supplementary Figure 3.

Second, we found that Phen2Disease outperformed Phen2Disease-double significantly in the real data. For the simulated data cohorts, Phen2Disease and Phen2Disease-double had a similar performance. For example, for the simulated data Cohort 2 (N=1000), Phen2Disease and Phen2Disease-double ranked 44.20% and 41.30% of the real disease-causing genes in Top1, respectively. The MRR of Phen2Disease and Phen2Dise-double were 56.12 and 54.74, respectively. In contrast, for the real data cohorts, the performance of Phen2Disease was significantly better than that of Phen2Disease-double. This also indicates that our integrated model Phen2Disease demonstrates strong robustness to different data cohorts.

### Heterogeneity of Data Cohorts

To further examine the prediction differences across the six real data cohorts used in our experiments, we conducted a heterogeneity analysis of all the data cohorts. As such, we first combined all the patients from the data cohorts. Our Phen2Disease produced a prediction score for each patient with the causative gene and a corresponding priority (rank). We plotted the scores and ranks in a scatter plot for all the patients as Figure 6A. From this figure, the ranking of cases shows a significant downward trend, as the Phen2Disease prediction score increases. This indicates that for all patients, the higher the prediction score, the higher the priority ranking of the correct gene will be.

**Figure 5.**
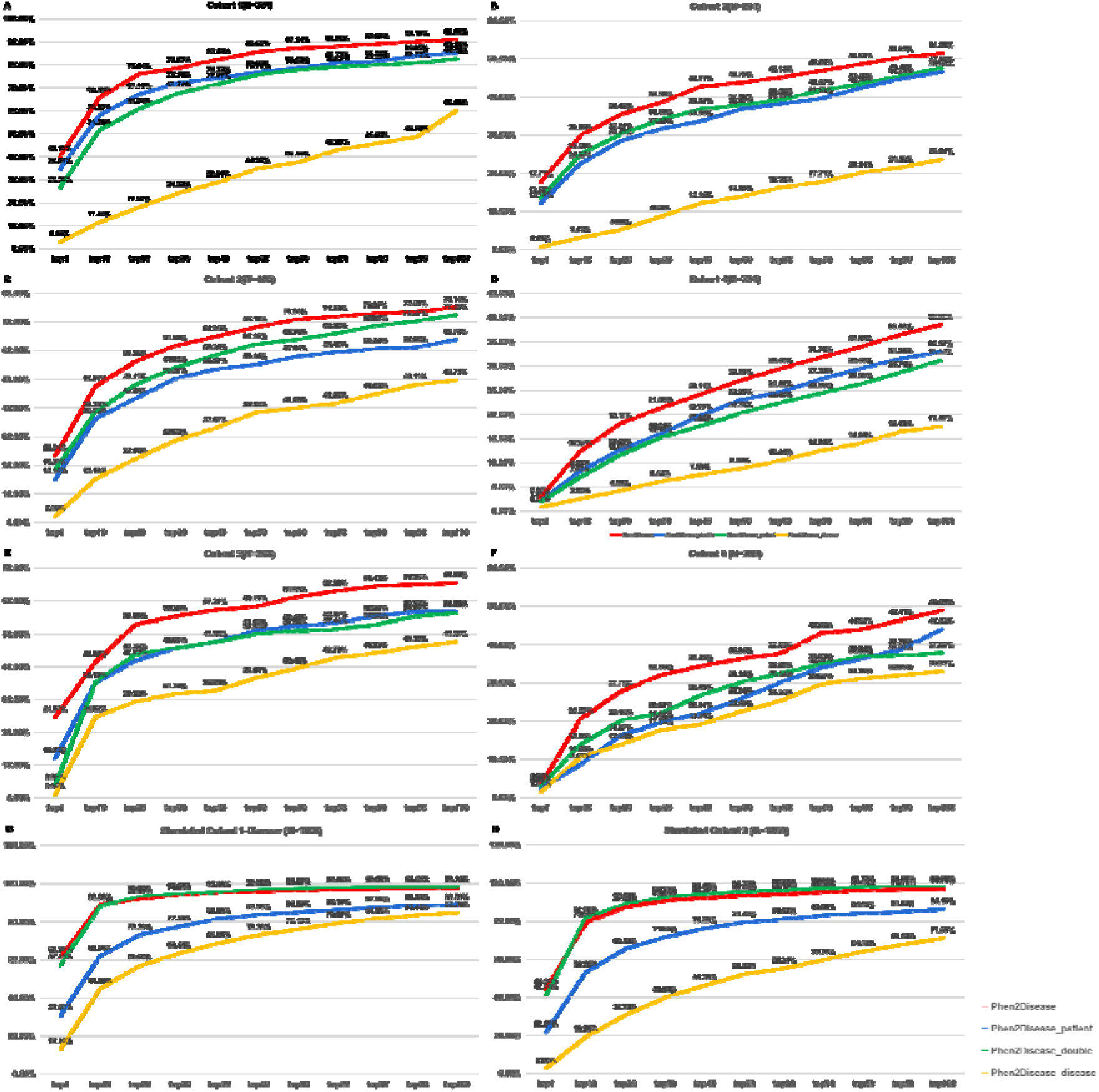
Comparisons of Phen2disease with its three single variants of Phen2disease_patient, Phen2disease-double and Phen2disease-disease in Cohort 1 (384 cases, Figure 5A), Cohort 2 (281 cases, Figure 5B), Cohort 3 (185 cases, Figure 5C), Cohort 4 (784 cases, Figure 5D), Cohort 5 (208 cases, Figure 5E), Cohort 5 (209 cases, Figure 5F), simulated Cohort 1 (1000 cases, Figure 5G), and simulated Cohort 2 (1000 cases, Figure 5H) in gene and disease ranking results.

**Figure 6.**
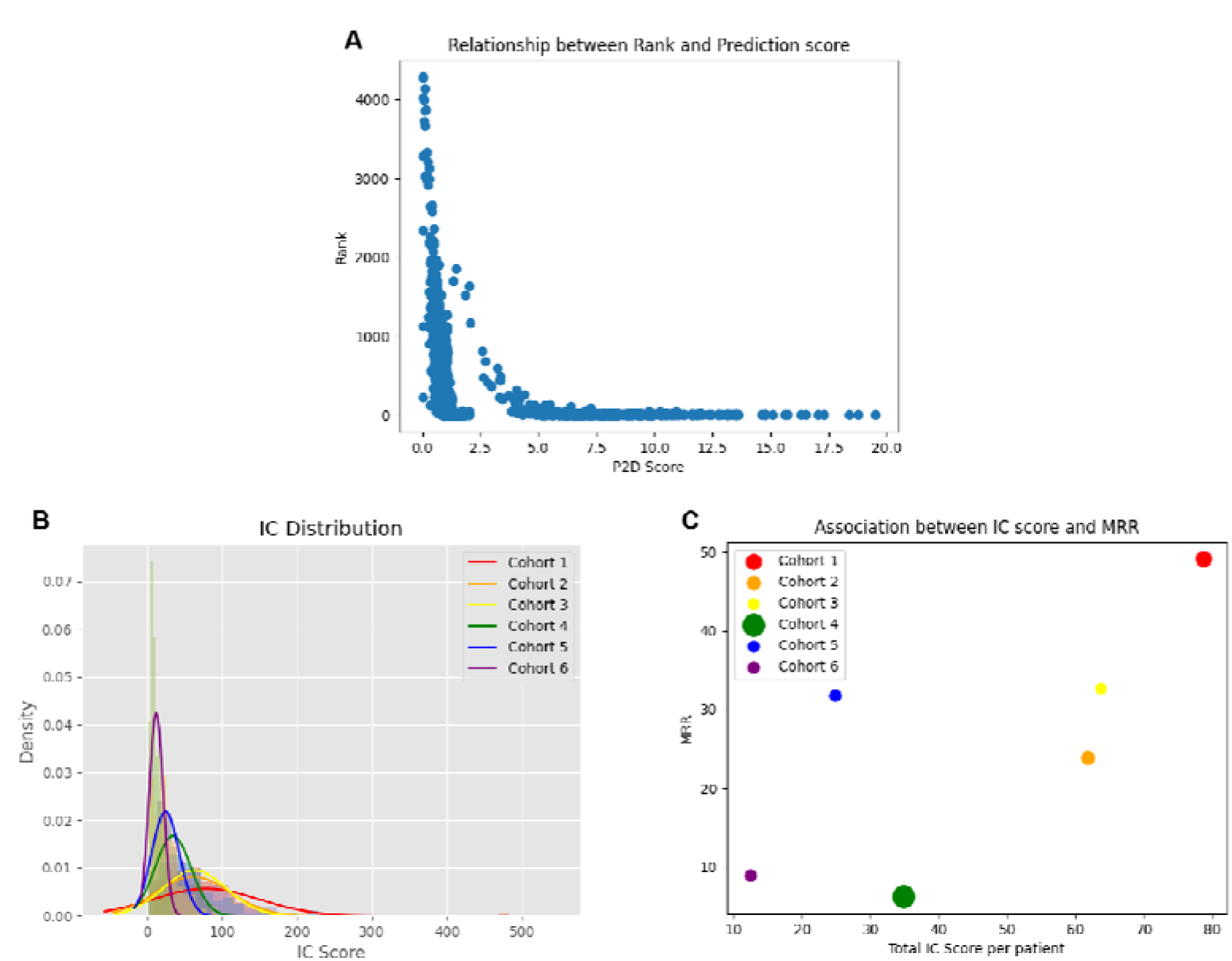
Correlation analysis of data cohort characteristics and prediction results (“P2D Score means Phen2Disease predicted scores). **A.** A scatter plot of true genetic ranking versus Phen2Disease predicted scores for all patients in all real data cohorts. **B.** Distributions of IC scores for six real data cohorts. **C.** A scatter plot of the relationship between total IC score per patient and MRR for six data cohorts (where the size of the circle corresponds to the number of cohort cases and Spearman correlation=0.7143).

Second, we analyzed each data cohort by separately plotting its distribution of IC scores for the six data cohorts in Figure 6B. The IC scores of each cohort were obtained by summing the IC scores of each patient using the filtered set of patient phenotypes, and finally averaging the IC scores of the patients in the data cohorts. We found that generally the larger the average IC scores, the corresponding data cohorts would be more predictable in terms of achieving higher values of the MRR metric. For example, Cohort 1 has a large average IC score, and its corresponding MRR value of 50.25 is relatively high. Conversely, the smaller the average IC score, the more difficult to predict the data cohort. For example, the average IC score of Cohort 6 is small, and its corresponding MRR value of 10.48 is also relatively low. The reason for this may be that data cohorts with high IC scores correspond to disease phenotypes with more specific information, so it is easier to match the corresponding disease and find the causative gene. In Figure 6C, we plotted the IC scores and the corresponding MRR for each data cohort as a scatter plot. Overall, a strong positive correlation is clearly shown, with a Spearman correlation coefficient of 0.7143. This indicates that to some degree, the IC score can reflect the prediction difficulty of a data cohort to some extent. However, we found that Cohort 5 was exceptional, with a relatively high MRR of 30.84, but a relatively low average IC score. Therefore, we carefully analyzed this data cohort with the following results. The average number of phenotypes in Cohort 5 was only 4.79, so the overall average IC score was relatively low. The patients in Cohort 5 were composed of 208 Chinese Han Chinese with a genetic diagnosis of skeletal disorders from 2009 to 2019, uniformly for skeletal genetic disorders. As the homogeneity of the data in Cohort 5 is higher, the patients with the same type of disease genes are relatively easy to predict, resulting in a better overall prediction (see Supplementary Table 17).

Finally, we consider each of the five independent data cohorts of data cohort 2 and plot scatters of the IC scores and corresponding MRRs for each of the ten independent data cohorts. We obtain a spearman correlation of 0.6242, which is still consistent with our conclusion. The detailed information can be found in Supplementary Figures 4-5.

### Case study: Phen2Disease helps physicians identify important phenotypes corresponding to diseases and genes

We provide a case study to further validate the performance of Phen2Disease in this section. Since Phen2Disease and Phrank have a similar overall process, we compare them in this case study. We use a real case from our Cohort 1: “Bluteau-2018-SAMD9L-UB085”. The corresponding disease and causative genes are “MYELOCEREBELLAR DISORDER (OMIM: 159550)” and “SAMD9L”, respectively. We compared the priority ranking of the different methods. For gene prioritization, Phen2Disease accurately ranked the SAMD9L gene in 1st place, followed by Phen2Disease-patient 10th, Phrank 23th and Phen2Disease-disease 54th.

Phen2Disease calculates a maximum matching score for each phenotype of the patient and the disease. As shown in Figure 7, Phen2Disease assigns scores to the patient and disease phenotypes according to the maximum matching score. There are three common phenotypes to both the patient phenotype and the disease phenotype: “HP:0001876” (Pancytopenia), “HP:0002317” (Unsteady gait) and “HP:0002500” (Abnormal cerebral white matter morphology). As shown in Figures 7A and 7B, we ranked these three phenotypes high in both patient and disease phenotypes, with phenotypes “HP:0001876” (Pancytopenia), “HP:0002317” (Unsteady gait) ranking in the top two. The third-ranked patient phenotype “HP:0001321” (Cerebellar hypoplasia) scored 3.10. It belongs to the same non-sub-ontological node phenotype “HP:0000707” (Abnormality of the nervous system) as the majority of the corresponding disease phenotypes. Moreover, the phenotype “HP:0001321” (Cerebellar hypoplasia) and the disease phenotype “HP:0001272” (Cerebellar atrophy) share a deep kinship node “HP:0011283” (Abnormality of the metencephalon). This suggests that this phenotype “HP:0001321” (Cerebellar hypoplasia) is associated strongly with the disease of MYELOCEREBELLAR DISORDER. Similarly, we used Phrank to calculate scores for each phenotype as well and ranked the scores from the highest to lowest as shown in Figure 7C. Phrank ranked only “HP:0002317” (Unsteady gait) in the first position, while the other two important overlapping phenotypes “HP:0001876” (Pancytopenia) and “HP:0002500” (Abnormal cerebral white matter morphology) were ranked outside the top 10 (the rank of these two phenotypes were 12th and 19th, respectively). Phrank did not correctly identify this disease, either.

**Figure 7.**
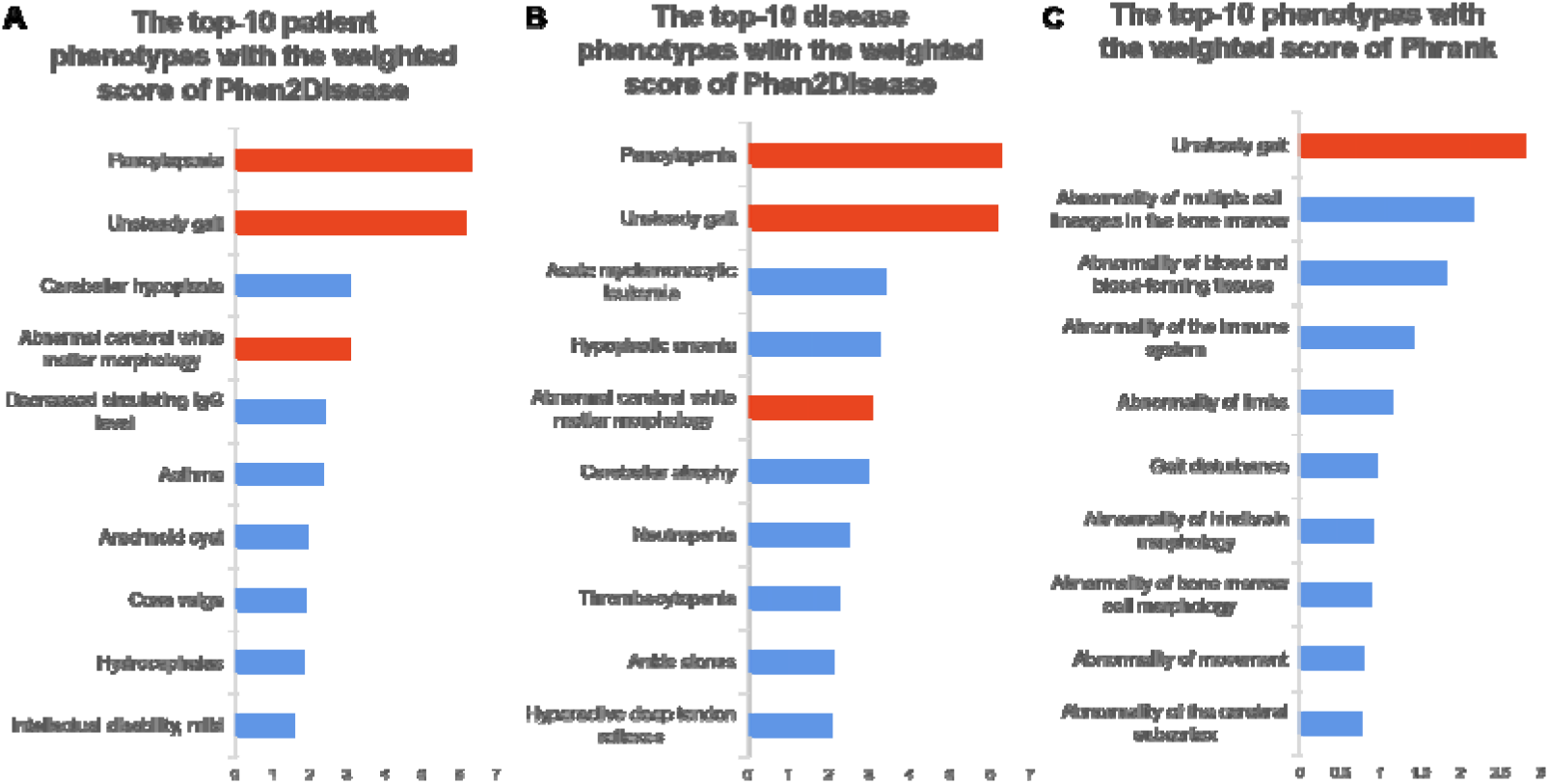
The top-10 phenotypes with weighted scores of Phen2Disease and Phrank (Red represents phenotypes shared by patient phenotypes and disease phenotypes). **A.** The top-10 patient phenotypes with the weighted score of Phen2Disease, where the weighted score is the maximum matching score calculated by phen2disease for each patient’s phenotype. **B.** The top-10 disease phenotypes with the weighted score of Phen2Disease, where the weighted score is the maximum matching score calculated by phen2disease for each disease’s phenotype. **C.** The top 10 phenotypes with a weighted score of Phrank, where the weighted score is the matched IC score calculated by Phrank.

Finally, we censored each of the phenotypes for the current patient and then performed gene prioritization ranking. We focused on three common phenotypes of the patient and disease, HP:0001876 (Pancytopenia), HP:0002317 (Unsteady gait) and HP:0002500 (Abnormal cerebral white matter morphology)), as shown in Table 4. It shows that the higher the IC score of a phenotype, the lower the causative gene will be ranked. For example, if the phenotype HP:0001876 (Pancytopenia) with an IC score of 6.29 is removed, the disease-causing gene prioritization ranking changes from the first to the seventh.

**Table 4.**
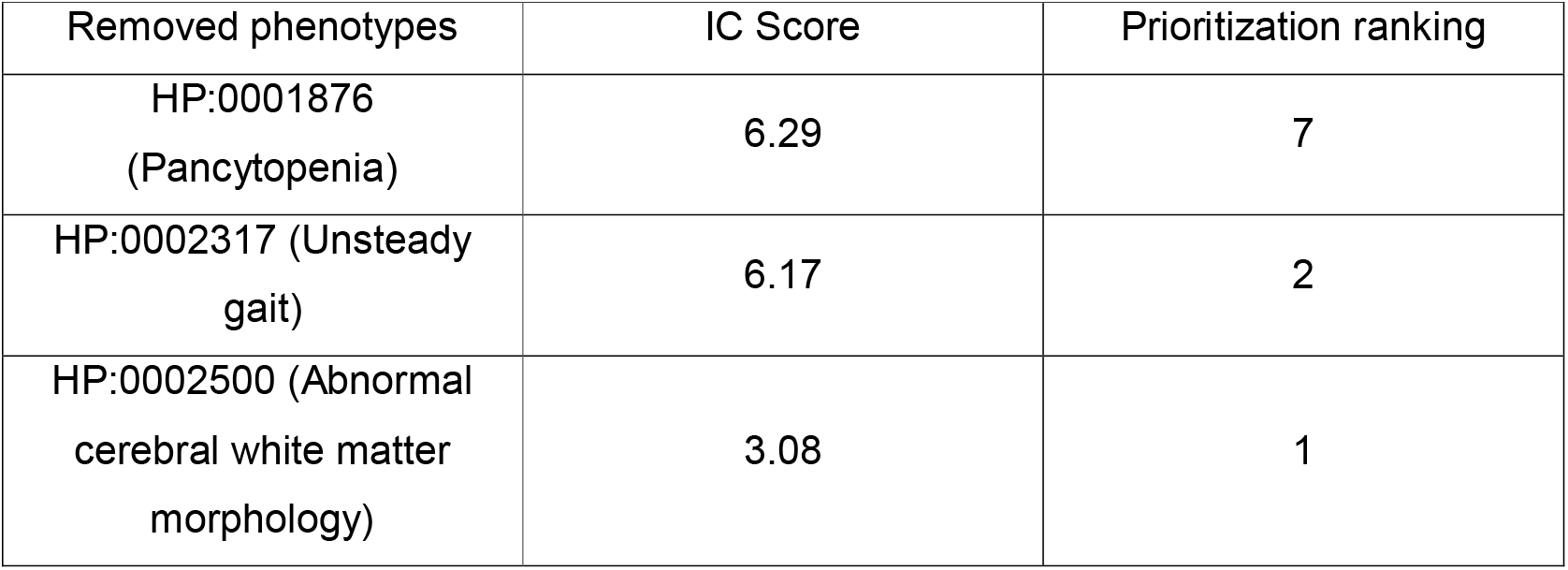
Prioritization by Phen2Disease-double of disease-causing genes after removal of a patient phenotype.

## DISCUSSION

Phen2Disease can prioritize disease-causing genes to improve the speed of diagnosis so as to reduce the burden on clinicians. The HPO structure information and IC weights are used to calculate the similarity score between patient and disease phenotypes. In particular, IC weights are associated with different term nodes to measure the degree of their contributions to the similarity. From the experiments, we found that patient-centric and patient-disease-centric scores that are the similarity between medical subject terms have a good prioritization ability. In our final model, we combined these two scores to enable more effective prioritization of a patient’s likely disease and causative gene from his/her phenotypes.

Overall, Phen2Disease performs the best in the six real data cohorts and two simulated cohorts. It outperforms the BASE_IC method, which directly considers the intersection IC of the patient phenotypes and the disease (gene) phenotypes. In addition, we have thoroughly compared Phen2Disease with the baseline methods of Phenolyzer, Phrank, Phen2Gene, LIRICAL, PhenoApt, and AMELIE against the same patient phenotypes. All these methods have certain limitations. For example, PhenoApt performs poorly in Cohort 1 and Cohort 2, which are not used in Chen et al. [29]. Differing from Phen2Disease, Phrank uses only the local information of its parent node to calculate the information content of a node. Specifically, it uses the intersection of phenotype sets to calculate the similarity of two phenotype sets, which is too coarse-grained. This has been validated by the results from the two real cases reported in Supplementary Table 18, showing that Phrank ranked the patients greater than 40th for both true causative genes and diseases. In contrast, our integrated model of Phen2Disease can accurately identify the true disease-causing genes and diseases.

AMELIE performed the best in two data cohorts (Cohort 2 and Cohort 3) in terms of Top1 hits, both of which are mainly curated from the literature. The performance of AMELIE will be significantly reduced if no known related literature exists. For example, we generated the simulated Cohort 2 using the simulated genetic phenotyping approach of Chen et al [30]. In these previously unreported cases, the prioritization ability of AMELIE was substantially reduced. On the contrary, our model achieves the best prioritization results in the last two real data cohorts and the simulated ones. It shows that our model still shows the very good ability of disease gene prioritization with good robustness in the presence of polycentric data and interspersed with different noises (the robustness of our model to different data cohorts can be also demonstrated in Supplementary Figures 6-13). In addition, with very few HPO terms, AMELIE loses its prioritization capability. For example, in Cohort 5, AMELIE performs poorly in terms of the TopK and MRR metrics. This is because when the number of terms is too small, AMELIE lists too many candidate genes in the same prioritization level. For one case of “DISCO-AX120” in Cohort 5 with only one HPO term “HP:0004322”, AMELIE scored 99.99 for 2334 candidate genes. Among them, the correct causative gene was “FBN1” with scoring the same 99.99, leaving no way of ranking genes for screening. However, our method produced a score of 1.2812 for the gene “FBN1”, ranked as the fourteenth. The reason for this is that our Phen2Disease uses both patient and disease phenotypes. The same patient phenotypes can further be distinguished in Phen2Disease by ranking the degrees of their associations with different diseases. The final Phen2Disease score integrates two normalized scores from the patient and diseases. Even if a patient has only one single patient HPO term, our scores are still different as a result of measuring the various degrees of its association with all disease genes. Therefore, Phen2Disease has a good performance on prioritization.

Phen2Disease improves its prioritization performance by using integrated information. Specifically, we find that the ranking results are more effective by using an integration score than by using the patient-centric and (or) disease-centric ones. The reasons for this are as follows. First, diagnoses based on patient phenotypes will be more accurate and straightforward than those based on disease phenotypes. A disease phenotype contains a variety of patient symptoms, but these symptoms are not present in all patients. This means that a patient’s phenotype may occupy only a fraction of all phenotypes of the corresponding disease. Therefore, we used a scoring method based on a patient phenotype-centric rather than on a disease-centric disease genetic prioritization. Second, there is also a drawback if a diagnosis is based only on the patient’s phenotype. This is because patients describe their disease pain differently even if they have the same disease, for example. In addition, the patient’s description may be inaccurate or miss some important symptoms. Therefore, we compute a combined patient-disease-centric score by using the information from both a patient and disease. Finally, we integrate the two similarity scores for diagnosis in the final model. The results in Figure 5 show that the integrated model is indeed superior in the clinical diagnosis of causative genes and diseases.

As reported before, our data cohorts have a high degree of heterogeneity. This is because they came from different data sources and different collection methods (Table 2). First, some data cases were from the published medical literature. In particular, the data Cohort 2 has 14 cases with a unique disease-causing gene (TAF1), from an article in the American Journal of Human Genetics [34]. Second, some data cases were extracted from medical reports using natural language processing (NLP) methods. For example, data cohort 2 had 72 cases from 61 molecular case study articles from Cold Spring Harbor. All the cases were first obtained using the Aho-Corasick algorithm embedded in Doc2HPO [26] and were then reviewed to remove negative and duplicate terms. Finally, data cases were also taken from real patients in hospitals and manually extracted and screened by specialized physicians. For example, Cohort 6 was extracted from 209 in-house cases at the CKCCL and was phenotypically screened by a dedicated certified genetic analyst [45]. Partially because of the diversities of these data, the compared methods performed differently across the data cohorts. However, our method Phen2Disease consistently performed well on all data cohorts. From the perspective of the individual data record of patients, we further found that patients with high IC scores had more specific information and their predictions would be more accurate. In this sense, the average IC score can somewhat be indicative of the difficulty of prediction for the data clusters. Also, other factors can influence the prediction results, such as the high homogeneity of the data cohort, which would lead to a better prediction.

Except for good performance, our Phen2Disease is of high clinical relevance. First, Phen2Disease can accurately prioritize both candidate diseases and genes, as well as highlight the most important patient and disease phenotypes. As such, it will be helpful for physicians to diagnose diseases and screen for causative genes. Second, it is convenient for physicians to use Phen2Disease in practice from its design perspective. Phen2Disease relies only on the widely used open public databases of HPO and HPO-A to produce accurate disease gene prioritization. The code of Phen2Disease is open source. Finally, Phen2Disease is highly interpretable by fully utilizing the information content of the hierarchical HPO DAG nodes to calculate similarity scores.

Phen2Disease also has some limitations. First, Phen2Disease prioritizes candidate diseases before prioritizing candidate disease-causing genes. Therefore, if the correct disease is not annotated by the current database, it is not possible to effectively match the correct disease and the causative gene. Second, Phen2Disease relies on the correct matching between patient and disease phenotypes. However, different institutions have different criteria for collecting patient phenotypes. An incomplete match between patients’ clinical non-specific phenotypes and disease phenotypes can lead to biased prioritization results. With a complete knowledge base or more accurate descriptions of patient phenotypes, the performance of Phen2Disease can be further improved to enhance clinical efficiency. Third, the current experimental results demonstrate that Phen2Disease has a good ability of disease gene prioritization in skeletal-type patients (Cohort 5), but this needs more data for further study and validation. Overall, Phen2Disease performs well for patients with different types of diseases with strong robustness and generalizability. In the future, we will conduct more experiments on Phen2Disease with data from different categories of diseases.

## ACKNOWLEDGEMENT

We thank Pro. Kai Wang (University of Pennsylvania) for the suggestions for adding the data cohort 6, analyzing the heterogeneity of data cohorts and insightful comments on the manuscript.

